# Comparing global and regional maps of intactness in the boreal region of North America: Implications for conservation planning in one of the world’s remaining wilderness areas

**DOI:** 10.1101/2020.11.13.382101

**Authors:** Pierre Vernier, Shawn Leroux, Steve Cumming, Kim Lisgo, Alberto Suarez Esteban, Meg Krawchuck, Fiona Schmiegelow

## Abstract

Though North America’s boreal forest contains some of the largest remaining intact and wild ecosystems in the world, human activities are systematically reducing its extent. Consequently, forest intactness and human influence maps are increasingly used for monitoring and conservation planning in the boreal region. We compare eight forest intactness and human impact maps to provide a multi-model assessment of intactness in the boreal region. All maps are global in extent except for Global Forest Watch Canada’s Human Access (2000) and Intact Forest Landscapes (2000, 2013) maps, although some global maps are restricted to areas that were at least 20% treed. As a function of each map’s spatial coverage in North America, the area identified as intact ranged from 55% to 79% in Canada and from 32% to 96% in Alaska. Likewise, the similarity between pairs of datasets in the Canadian boreal ranged from 0.58 to 0.86 on a scale of 0-1. In total, 45% of the region was identified as intact by the seven most recent datasets. There was also variation in the ability of the datasets to account for anthropogenic disturbances that are increasingly common in the boreal region, such as those associated with resource extraction. In comparison to a recently developed high resolution regional disturbance dataset, the four human influence datasets (Human Footprint, Global Human Modification, Large Intact Areas, and Anthropogenic Biomes), in particular, omitted 59-85% of all linear disturbances and 54-89% of all polygonal disturbances. In contrast, the global IFL, Canadian IFL, and Human Access maps omitted 2-7% of linear disturbances and 0.1-5% of polygonal disturbances. Several differences in map characteristics, including input datasets and methods used to develop the maps may help explain these differences. Ultimately, the decision on which dataset to use will depend on the objectives of each specific conservation planning project, but we recommend using datasets that 1) incorporate regional anthropogenic activities, 2) are updated regularly, 3) provide detailed information of the methods and input data used, and 4) can be replicated and adapted for local use. This is especially important in landscapes that are undergoing rapid change due to development, such as the boreal forest of North America.

## Introduction

North America’s boreal forest contains some of the largest remaining intact ecosystems in the world (Potapov et al. 2017, Watson et al. 2018). However, the rapid expansion of industrial activities such as forestry, mining, and oil and gas exploration into increasingly accessible landscapes is systematically reducing its extent (Bradshaw et al. 2009, CEC 2010, Schindler and Lee 2010, Brandt et al. 2013, Venier et al. 2014). Large intact areas support biodiversity, ecological and evolutionary processes including wildlife migrations and natural disturbances, and ecosystem services such as carbon capture and sequestration (Mittermeier et al. 2003, Leroux et al. 2010, Watson et al. 2016). They also play an important role in climate change mitigation (Price et al. 2013, Melillo et al. 2016, Carroll and Noss 2020) and can serve as ecological baselines to guide sustainable land management practices (Arcese and Sinclair 2016). Despite their importance and recent calls for the expansion of protected areas in intact or wilderness regions (Betts et al. 2017, Dinerstein et al. 2017, Tilman et al. 2017), the global erosion of wilderness areas has exceeded their rate of protection (Watson et al. 2016). To identify and conserve additional wilderness and intact areas, reliable and up-to-date spatial information is required. This has led to the production of several global and regional datasets that attempt to map anthropogenic disturbances or their complement, areas with little or no evidence of human activities (McCloskey and Spalding 1989, Bryant 1997, Sanderson et al. 2002, Potapov et al. 2008b, Hansen et al. 2013). The maps vary in methodology, spatial and temporal characteristics, and most importantly, the area estimated to be intact in the boreal region. Consequently, a comparison of map products would assist conservation planners and researchers with the selection of the most appropriate product(s) for their purposes.

The boreal region of North America covers 6.3 million km^2^, of which 88% is in Canada and 12% is in Alaska (Brandt et al. 2013). In Canada, 11.4% of the region is currently under some form of protection (CPCAD 2019), and both globally and in Canada, there is increasing recognition of the need to expand protected areas while opportunities remain. In response, the United Nations Convention on Biological Diversity developed a set of goals (“Aichi Targets”) for protecting biodiversity which includes a target of 17% of terrestrial areas conserved by 2020 (Butchart et al. 2016), with a proposed increase to 30% by 2030 (Dinerstein et al. 2019). At the regional level, the Governments of Ontario and Quebec have committed to setting aside 50% of the boreal region of each province in various levels of protection in anticipation of future resource development (Government of Quebec (Minister of Natural Resources and Wildlife) 2009, Hansen et al. 2010, OMNR 2013). Beyond their use in conservation planning, intact forests are being considered as a policy instrument in forest conservation and management and have recently been integrated into the certification standards of the Forest Stewardship Council (FSC 2015). The forests and peatlands of the boreal region also are important carbon sinks and the maintenance of intact forest landscapes may be part of a natural solution to carbon sequestration and CO_2_ reduction (Griscom et al. 2017). Consequently, intact areas have an important role to play in protected area design and as control areas against which the impacts of human activities on biodiversity can be compared within an adaptive management framework (Lindenmayer et al. 2006, Watson et al. 2009).

In the boreal region where forests dominate the landscape, wilderness or intact areas have much in common with the concept of the Intact Forest Landscape (IFL), defined as “a seamless mosaic of forest and naturally treeless ecosystems within the zone of current forest extent, which exhibit no remotely detected signs of human activity or habitat fragmentation and is large enough to maintain all native biological diversity, including viable populations of wide-ranging species” (Potapov et al. 2008). Depending on the IFL dataset, intact areas must be a minimum of 1,000 ha to 50,000 ha (Potapov et al. 2008, Lee et al. 2010). An area becomes non-intact through the accumulation of human impacts, often related to resource extraction activities such as logging, mining, oil and gas development, and their associated roads. In this context, intactness is considered to be a structural descriptor of landscapes that reflects the absence of anthropogenic disturbances as measured from thematic (e.g., roads) and remote sensing data.

Several global and regional initiatives have attempted to map the overall condition of the world’s ecosystems in the past 30 years (Table 1). The initiatives can be divided into two broad groups based on their objective: intactness mapping and human influence mapping. The intactness mapping approach attempts to map remaining areas with little or no human activities by removing anthropogenic disturbances that are detectable using satellite imagery and other input data, and sometimes applying a zone of influence buffer. Resulting areas are considered free from significant human pressures. In contrast, the human influence mapping approach combines multiple disturbance layers into an overall map showing areas of low to high disturbances. Areas with the least amount of disturbance can then be reclassified to identify relatively intact areas. Among the intactness mapping approaches, the World Wilderness Areas map was one of the first global initiatives (McCloskey and Spalding 1989). To qualify, areas had to be ≥ 4,000 km^2^ after eliminating all areas within 6 km of human infrastructures e.g., roads and settlements. Subsequently, the Frontier Forests initiative, produced by the World Resources Institute, also attempted to map the world’s remaining large intact natural forests (Bryant 1997). No explicit minimum size was specified and, similar to wilderness areas, human disturbances due to traditional activities were considered acceptable. The concept, however, has been questioned for its utility for identifying high priority conservation areas because of the methods and criteria used to define intact forests (Innes and Er 2002). More recently, Global Forest Watch (GFW) built upon the concept of the Frontier Forest to develop a global map of intact forest landscapes (IFLs), delineated using specific criteria related to minimum size, patch width, and corridor width (Potapov et al. 2008, 2017). The global IFL maps were produced for the years 2000, 2013 and 2016. The approach has also been applied at a regional scale in Canada for the years 2000 and 2013 by GFW Canada (Lee et al. 2010, Smith and Cheng 2016), although it diverges from the global definition with respect to some of the criteria, for example the treatment of wildfires (Lee 2009).

**Table 1.** General characteristics of intactness and human influence maps reviewed in this study. Input data sources and methodological characteristics of intactness and human impact maps reviewed in this study. References and links to all datasets are provided in the supporting information (S1_datasets.md). Note that the Canada Human Access dataset is listed under intactness type maps since it was used as an input to the Canada IFL maps and both were produced by GFWC.

| Dataset | Years<sup>1</sup> | Geographic extent | Format | Scale / Resolution<sup>2</sup> | Measurement scale | Buffer distance<sup>3</sup> | Minimum patch size of intact area | Reclass<sup>4</sup> | Thematic maps<sup>5</sup> | Satellite imagery |
| --- | --- | --- | --- | --- | --- | --- | --- | --- | --- | --- |
| **INTACTNESS TYPE MAPS** |  |  |  |  |  |  |  |  |  |  |
| Frontier forests<sup>8</sup> (FF) | 1996 | Global; terrestrial ecosystems | Vector | 1:8,000,000 (~16 km<sup>2</sup>) | Binary; frontier or not frontier, with threat levels | n/a | Generally, > 50,000 ha | n/a | World Forest Map<sup>9</sup> and Wilderness Areas map (McCloskey and Spalding 1989) used by > 90 experts to define large forested areas free of roads, settlements, etc. | No |
| Canada human access (HA) | 2010 | Canada; terrestrial ecosystems | Vector | 1:1,000,000 (~0.25 km<sup>2</sup>) | Binary; human access or not | 0.5 km | n/a | No human access = intact | Roads, mines, clearcuts, wellsites, pipelines, transmission lines, and agricultural clearings | anthropogenic disturbance layers (S2) |
| Boreal ecosystem anthropogenic disturbance (BEAD) | 2010, 2015 | Canada boreal; 51 caribou ranges<sup>10</sup> | Vector | 1:50,000 | Binary; human access or not | unbuffered | n/a | No human access = intact | Hydro reservoirs (GFWC) | Landsat 5 (30-m; 2008-2010) and Landsat 8 (15- and 30-m; 2015-2017) |
| Canada intact forest landscapes (CIFL) | 2000, 2013 | Canadian Boreal; 11 forested ecozones | Vector | 1:1,000,000 (~0.25 km<sup>2</sup>) | Binary; intact or not intact | 1 km around highways; 0.5 km around other disturbance types | 5,000 ha boreal & taiga ecozones; 1,000 ha temperate ecozones, which occur along southern edge of Brandt's boreal | n/a | Linear features (roads, cutlines, etc.), reservoirs, settlements; HA2010 and BEAD2010 | Landsat 5 & 7 (1988-2006; 28.5m); Landsat composite (~2013; 30m); anthropogenic disturbance layers and forest disturbance dataset (S2) |
| Global intact forest landscapes<sup>6</sup> (GIFL) | 2000, 2013, 2016 | Global; forested zones - tree canopy >20% | Vector | 1:1,000,000 (~0.25 km<sup>2</sup>) | Binary - intact or not intact | 1 km | 50,000 ha, at least 10-km wide at broadest place, at | n/a | Roads, settlements, scanned topographic maps | Landsat 5 (~1990; 30m) and Landsat 7 (~2000; 30m); MODIS VCF 2000 |
|  |  | & area >4 km<sup>2</sup> (MODIS 2000) |  |  |  |  | least 2-km wide in corridors |  |  | (percent tree cover; 0.5 km<sup>2</sup>); Landsat composite (~2013; 30m) |
| **HUMAN INFLUENCE TYPE MAPS** |  |  |  |  |  |  |  |  |  |  |
| Human footprint<sup>7</sup> (HFP) | 2000, 2005, 2010, 2013 | Global; terrestrial ecosystems; stratified by biomes & ecoregions | Raster | 1 km<sup>2</sup> | Ordinal; 0-50 (low to high); sum of ranks of human pressures | Two influence zones: 0-2 km & 2-15 km | 50,000 ha | HF1993 = 0; HF2009 = 0 | 7 anthropogenic stressors (human population density, built-up area, cropland, livestock, forest cover change, roads, night-time lights) | Various including global land cover (GLC2000; 1 km<sup>2</sup>) and GlobCover 2009 (300m) |
| Anthropogenic biomes (Anthromes, AB) | 2000, 2005, 2010, 2015 | Global; terrestrial ecosystems | Raster | ~5 km<sup>2</sup> | Categorical; 6 groups, 19 classes | n/a | n/a |  | 6 anthropogenic stressors (human population density, built-up area, cropland, rice area, irrigated area, pasture) | MODIS VCF 2000 (Percent Tree Cover; 0.5 km<sup>2</sup>) |
| Global human modification (GHM) | 2016 | Global; terrestrial ecosystems | Raster | 1 km<sup>2</sup> | Continuous 0-1 (low to high); proportion of landscape modified | n/a | n/a |  | 13 anthropogenic stressors (human population density, built-up area, cropland, livestock, major roads, minor roads, two-tracks, railroads, mines, oil wells, wind turbines, power lines, night-time lights) |  |
| Very low impact areas (VLIA) | 2015 | Global; terrestrial ecosystems | Raster | 1 km<sup>2</sup> | Binary; 2 classes |  |  | n/a | 7 anthropogenic stressors (human population density, built-up area, cropland, livestock, forest cover change, roads, night-time lights) |  |

<sup>1</sup> If the year of the dataset is not provided, we use the date of latest imagery using as input.
<sup>2</sup> Values in brackets for vector maps indicate approximate effective grid resolution, similar to minimum mapping unit for polygonal data.
<sup>3</sup> The distance around disturbances that is removed from the estimation of intact areas.
<sup>4</sup> Map categories or values that were reclassified to indicate intactness.
<sup>5</sup> Human disturbances considered by the map producers; method of detection varied by map and disturbance type and included existing maps, satellite imagery and aerial photos.
<sup>6</sup> A key intermediate dataset was GFWC's Canada Access 2010 dataset, which was created as the initial step in creating the IFL maps.
https://globalforestwatch.ca/sites/gfwc/files/data/20140109B_Canada_Access_2010_metadata.html
<sup>7</sup> HFP1993 is an update to the original human footprint/human influence index dataset (circa 1993) [47]
<sup>8</sup> Frontier forests are large, ecologically intact, and relatively undisturbed natural forests [9]
<sup>9</sup> The World Conservation Monitoring Centre, The World Forest Map, (WCMC, Cambridge, 1996). <sup>10</sup> Partial updates to some caribou ranges in 2012.

Among the human influence mapping approaches, the best known is the Human Footprint (HFP), which provides a standardized measure of cumulative human pressures on the environment based on the extent of built environments (e.g., urban areas), crop land, pasture land, human population density, night-time lights, railways, roads and navigable waterways (Sanderson et al. 2002). The HFP has been updated several times while adhering to consistent methods, with the most recent dataset current to 2013 (Venter et al. 2016, Williams et al. 2020). All versions of the HFP can be reclassified to identify areas with little or no disturbances. Another well-known dataset is the Anthropogenic Biomes (Anthromes) map (Ellis and Ramankutty 2008) that classifies the terrestrial biosphere into 19 categories based on human interactions with ecosystems, including agriculture, urbanization, forestry and other land uses. It has also been updated for multiple years, with the most recent version current to 2015 (Ellis et al. 2020). For all versions, the most relevant categories for mapping intact areas are the wildland categories (wild forest, sparse trees and barren). Two additional recent datasets, the (Very) Low Impact Areas map (Jacobson et al. 2019) and the Global Human Modification map (Kennedy et al. 2019) also provide a cumulative measure of human modification of terrestrial lands across the globe at a 1-km^2^ resolution for the years 2015 and 2016, respectively. Both approaches are similar to the HFP approach but differ in the number and types of anthropogenic stressor datasets, as well as the methods used to calculate human influence (Riggio et al. 2020). At a regional scale, GFW Canada also developed the Human Access dataset for 2010 as an intermediary step to creating the Canada IFL 2013 map (Lee and Cheng 2014). The dataset maps recent linear and areal disturbances related to resource extraction but, unlike the IFL datasets, does not use a minimum size criterion, although buffers are applied around disturbances (Table 1).

The availability of an increasing number of intactness and human influence maps may lead to confusion about the suitability of the various products for conservation planning in the boreal region, especially since many of the datasets are global in extent. To identify the most suitable dataset, it would help to not only understand the differences in characteristics and assumptions of each map, but also how well their predictions agree with each other and against independent and higher-resolution regional data. A recent study comparing the four human influence maps at the global scale found that despite differences in methods and data, the datasets estimated similar percentages of low and very low human influence areas (Riggio et al. 2020). In the current study, we evaluated both intactness and human influence datasets (with the exception the World Wilderness Areas), focusing on the boreal region of Canada. Our overall objective was to compare and evaluate existing map products that can be used to measure intactness. Our specific objectives were to:

1. To compare intactness estimates across the boreal region of Canada and Alaska;
2. To quantify inter-map agreement and identify areas of common agreement; and
3. To evaluate the strengths and limitations of the maps to accurately identify anthropogenic disturbances common in the boreal region: oil and gas exploration, logging, roads, and mining.

We also provide two case studies in the supporting information: the first, on the effectiveness of intactness and human influence maps at identifying disturbances related to placer mining in west-central Yukon, and the second, on the sensitivity of intactness estimates to buffer size and minimum patch size in northern Alberta.

## Methods

Our overall study area comprises the spatial extent of the boreal and boreal alpine regions (i.e., the boreal region) of North America (Brandt 2009). However, most of our analysis is focused on a large subset of the Canadian boreal region (Figure 1, black outline), representing the intersection of the intactness and human influence maps evaluated. In total, we acquired eight freely available national and global maps that at a minimum, covered a large portion of the study region. All maps are global in extent except for those produced by GFW Canada which are restricted to Canada’s boreal and temperate forests. Four of the maps were produced for multiple periods for a total of 17 datasets (maps x years). Table 1 summarizes the general characteristics of each dataset including geographic extent, format, resolution, measurement scale, and source of input data. The table also provides information on mapping methods, including geographic extent, resolution, minimum size of intact areas, disturbance types, and buffering of disturbances.

**Figure 1.**
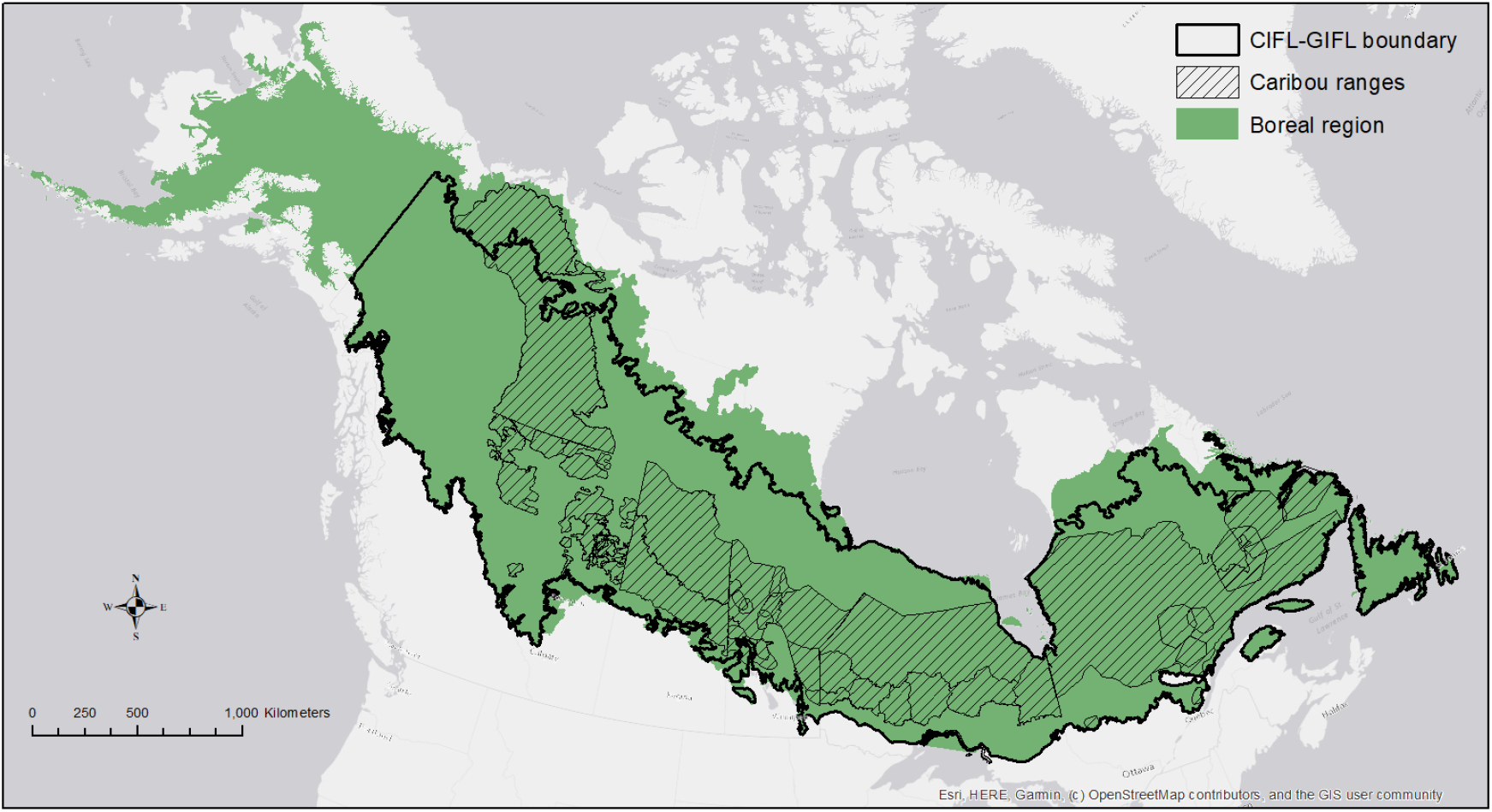
Extent of boreal region, including boreal and boreal alpine zones, in North America (Brandt 2009). The solid black line indicates the area of intersection of the eight intactness and human influence maps while the crosshatch pattern indicates the limits of the 51 caribou ranges that make up the 2015 BEAD dataset (Pasher et al. 2013).

### Intactness estimates and spatial agreement

The eight intactness and human influence map products varied in geographic coverage, mapped values, scale, coordinate system, and GIS file format. To enable comparisons of intactness estimates across maps across the boreal region, we converted all maps to an Albers Equal Area projection, clipped them to the boreal region of Canada and Alaska, and vectorized the raster datasets. We then reclassified the maps, where necessary, to binary “intactness” maps, with 1 indicating intact areas (i.e., little or no human influence) and 0 identifying non-intact areas. For the human footprint maps (HFP2000-13) and very low impact areas map (VLIA2015), all areas with little or no human influence (pixel values=0) were assigned a value of 1 while remaining areas were assigned a value of 0. For the global human modification map (GHM2016), we followed Riggio et al. (2002) and assigned pixels with values ranging from 0-0.01 a value of 1. For the HA2010 map, we eliminated all disturbance polygons from the boreal study region and assigned a value of 1 to the resultant areas. For the frontier forest map (FF), all polygons were assigned a value of 1 irrespective of their threat level. For the anthropogenic biome maps (AB2000-15), we assigned a value of 1 to the wildland categories i.e., wild forest, sparse trees and barren. Following map reclassifications, we calculated, for each map, the proportional area of the boreal region identified as intact. Since not all datasets covered the entire boreal region, the intact area identified by each dataset was divided by the total area of the map’s coverage within the boreal rather than by the area of the boreal. For Canada, we estimated intactness using all datasets. For Alaska, we used all datasets except for the GFW Canada datasets.

To evaluate the spatial agreement of datasets, we rasterized or resampled all maps to a 1000-m resolution, the most common resolution amongst the datasets evaluated. We restricted the spatial extent of the analysis to the area of intersection of all datasets. For the Canada IFL, Global IFL and HFP maps, we used the most recent maps. Frontier Forests was excluded because of its age and low original resolution. We quantified the area of spatial agreement (i.e., pairwise similarities) between the maps using Jaccard’s similarity coefficient (Fewster and Buckland 2001). The Jaccard coefficient measures the similarity between datasets, and is defined as the area of intersection of two datasets divided by the area of union. Values for the coefficient range from 0 (complete dissimilarity) to 1 (complete similarity). All analyses were conducted using R 4.0.2 (Team n.d.) and the sf (Pedzema 2018) and raster (Hijmans 2016) packages.

### Accuracy assessment

One of the main objectives of this study was to assess how well intactness and human influence datasets account for anthropogenic disturbances that are increasingly common in the boreal region, namely those associated with energy extraction, forestry, and mining. To do this, we used the boreal ecosystem anthropogenic disturbance (BEAD) dataset updated to 2015, a high-resolution dataset that was created specifically to identify disturbances in and around 51 boreal caribou ranges (Figure 1 hatched area; Pasher et al. 2013). Two broad types of disturbances were mapped within each range: 1) linear disturbances such as roads, seismic cutlines, and pipelines and 2) polygonal disturbances such as forest cutblocks, agricultural areas, and mining quarries. The dataset consists of both buffered and unbuffered linear and polygonal disturbances, covering 4.4 million km^2^ of the boreal encompassing 51 caribou ranges, and was produced using both 30- and 15-m resolution Landsat 8 data. In our analysis, we used the unbuffered dataset to evaluate the seven most recent datasets: HA2010, Canada IFL2013, Global IFL2016, HFP2013, AB2015, VLIA2015, and GHM2016. For each caribou range, we estimated of the proportion of linear and polygonal disturbance types omitted by each intactness and human influence map. These estimates are conservative because no buffer was applied to the disturbances.

## Results

### Intactness estimates and spatial agreement

All maps except for the Global IFL maps covered at least 98% of the boreal region of Canada as defined by Brandt (Brandt 2009) (Table 2, Supp Info S1). The three Global IFL maps covered 86% of the region. The total area identified as intact within the spatial extent of each map, ranged from 55-59% for the three global IFL maps (GIFL2000-16) to 89% for the four Anthromes maps (AB2000-15). The amount identified by the GHM2016, HA2010, and the four Human Footprint maps were very similar, ranging from 81-84%. In contrast, the FF1996 map was relatively low, with only 60% of the boreal identified as intact. The remaining datasets range between 71-76% for VLIA2015 and Canada IFL datasets (CIFL2000-13), respectively. Among multi-temporal datasets, the CIFL2013 map identified 3.6% less intact forest than the CIFL2000 map while the GIFL2016 map identified 3.4% less intact area than the GIFL2000 map. In contrast, the reduction in intact area between the newest and oldest HFP and AB maps was only 0.3 and 0.1%, respectively. In Alaska, the area identified as intact varied more widely than in Canada, ranging from 32% for the FF1996 map to 96% for the GHM2016 map (Table 2, Supp Info S1). The global IFL maps identified 19-21% more intact area in Alaska than in Canada with a 6% reduction between the oldest and newest maps. The Anthromes and Human Footprint maps had similar proportional values to Canada with little change over time.

**Table 2.** Comparison of the areal extent of dataset coverage within the boreal region and areas identified as being intact within each dataset. See Supplementary Information for distribution maps of each map in Canada and Alaska.

|  | Canada boreal region (5,519,764 km<sup>2</sup>) |  |  |  | Alaska boreal region (737,008 km<sup>2</sup>) |  |  |  |
| --- | --- | --- | --- | --- | --- | --- | --- | --- |
| Dataset | Dataset coverage (km<sup>2</sup>) | Coverage of boreal (%) | Intact area (km<sup>2</sup>) | Intact area (%) | Dataset coverage (km<sup>2</sup>) | Coverage of boreal (%) | Intact area (km<sup>2</sup>) | Intact area (%) |
| HA2010 | 5,519,764 | 100.0 | 4,565,591 | 82.7 |  |  |  |  |
| CIFL2000 | 5,394,980 | 97.7 | 4,029,533 | 74.7 |  |  |  |  |
| CIFL2013 | 5,394,980 | 97.7 | 3,837,668 | 71.1 |  |  |  |  |
| GIFL2000 | 4,746,030 | 86.0 | 2,780,919 | 58.6 | 475,765 | 64.6 | 379,475 | 79.8 |
| GIFL2013 | 4,746,030 | 86.0 | 2,652,463 | 55.9 | 475,765 | 64.6 | 355,012 | 74.6 |
| GIFL2016 | 4,746,030 | 86.0 | 2,619,094 | 55.2 | 475,765 | 64.6 | 350,983 | 73.8 |
| hfp2000 | 5,519,764 | 100.0 | 4,474,868 | 81.1 | 737,008 | 100.0 | 614,217 | 83.3 |
| hfp2005 | 5,519,764 | 100.0 | 4,481,331 | 81.2 | 737,008 | 100.0 | 614,444 | 83.4 |
| hfp2010 | 5,519,764 | 100.0 | 4,466,114 | 80.9 | 737,008 | 100.0 | 613,338 | 83.2 |
| hfp2013 | 5,519,764 | 100.0 | 4,464,139 | 80.9 | 737,008 | 100.0 | 613,685 | 83.3 |
| ab2000 | 5,519,764 | 100.0 | 4,919,781 | 89.1 | 737,008 | 100.0 | 645,668 | 87.6 |
| ab2005 | 5,519,764 | 100.0 | 4,919,588 | 89.1 | 737,008 | 100.0 | 645,668 | 87.6 |
| ab2010 | 5,519,764 | 100.0 | 4,921,931 | 89.2 | 737,008 | 100.0 | 645,668 | 87.6 |
| ab2015 | 5,519,764 | 100.0 | 4,918,939 | 89.1 | 737,008 | 100.0 | 645,427 | 87.6 |
| GHM2016 | 5,519,764 | 100.0 | 4,627,206 | 83.8 | 737,008 | 100.0 | 707,686 | 96.0 |
| VLIA2015 | 5,519,764 | 100.0 | 4,166,590 | 75.5 | 737,008 | 100.0 | 690,993 | 93.8 |
| FF1996 | 5,519,764 | 100.0 | 3,324,371 | 60.2 | 737,008 | 100.0 | 237,657 | 32.2 |

Spatial agreement, as measured by Jaccard’s similarity coefficient between intactness maps, ranged from a low of 0.58 between the GIFL2016 and VLIA2015 maps to a high of 0.86 between the AB2015 and GHM2016 maps (Table 3). Compared to the other maps, the similarity between the GIFL2016 map and all other maps except for CIFL2013 was 0.67 or less. With CIFL2013, it was 0.77. Most other paired comparison ranged between 0.70 and 0.85, indicating a relatively strong level of similarity. The intersection of the seven intactness maps revealed that 45% of the study region was identified as intact by all seven maps and an additional 18% by at least six of the maps (Figure 2). Only 3% of the study region was identified as not intact by all maps.

**Table 3.** Proportional agreement between each pair-wise map comparisons within Canada’s boreal region. Each entry represents the proportion of intact area in Map A (shown in the rows) that is also mapped as intact in Map B (shown in the columns). Comparisons were restricted to the area of intersection among the seven datasets (4,732,303 km^2^).

| Map | HA2010 | CIFL2013 | GIFL2016 | HFP2013 | AB2015 | GHM2016 | VLIA2015 |
| --- | --- | --- | --- | --- | --- | --- | --- |
| HA2010 | 1 |  |  |  |  |  |  |
| CIFL2013 | 0.85 | 1 |  |  |  |  |  |
| GIFL2016 | 0.67 | 0.77 | 1 |  |  |  |  |
| HFP2013 | 0.81 | 0.79 | 0.65 | 1 |  |  |  |
| AB2015 | 0.80 | 0.73 | 0.59 | 0.84 | 1 |  |  |
| GHM2016 | 0.81 | 0.77 | 0.62 | 0.83 | 0.86 | 1 |  |
| VLIA2015 | 0.74 | 0.70 | 0.58 | 0.75 | 0.77 | 0.84 | 1 |

**Figure 2.**
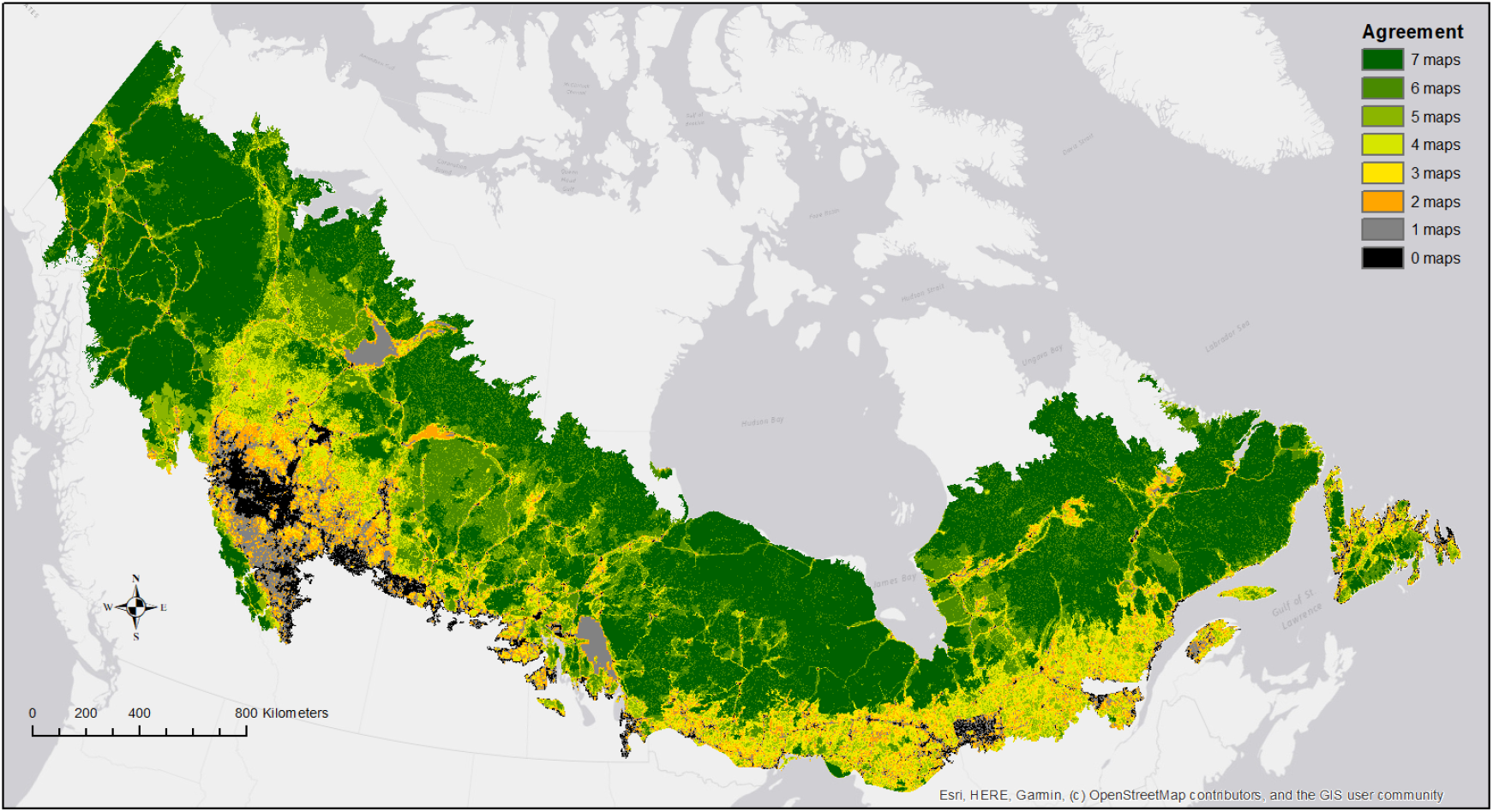
Spatial agreement between the seven most recent intactness (N=3) and human influence (N=4) maps. Dark green areas indicate areas identified as intact by all datasets. The percent area of agreement among: 7 maps = 45.5%, 6 maps=17.8%, 5 maps=8.8%, 4 maps=8.3%, 3 maps=6.6%, 2 maps=5.3%, 1 map=4.6%, 0 maps=3.0%.

### Accuracy assessment

The area identified as intact within the 51 caribou ranges by the seven intactness (N=3) and human influence (N=4) datasets ranged from 45% for the GIFL2016 map to 93% for the AB2015 map (Table 4). There were 31% more linear disturbances identified with the 15-m data than with the 30-m data. Most of the differences were due to seismic lines (41% more) and roads (22% more). In contrast, there was only 0.2% more polygonal disturbances identified with the 15-m data than with the 30-m data. Overall, the four human influence datasets (HFP2013, AB2015, VLIA2015, GHM2016), omitted 59-85% of all linear disturbances and 54-89% of all polygonal disturbances. In contrast, the intactness datasets (GIFL2016, CIFL2013, and HA2010) omitted between 2-7% of linear disturbances and 0.1-5% of polygonal disturbances.

**Table 4.**
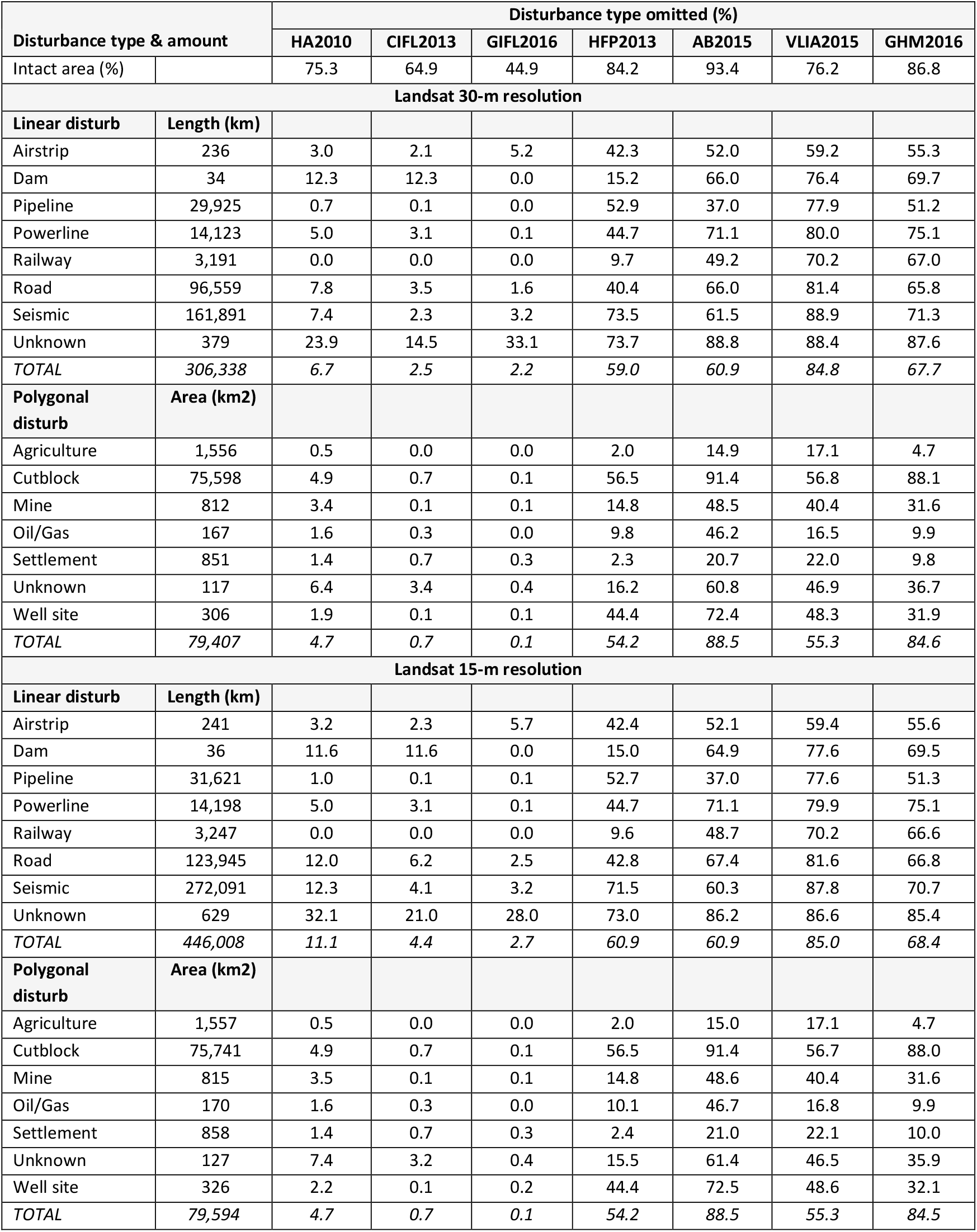
Percent length and area omitted (misclassified as intact) by each intactness and reclassified human influence map in the study area.

The most common linear anthropogenic disturbances in the study area were seismic cutlines, roads, pipelines and powerlines. Railways, airstrips, and dams also occurred, but to a much lesser extent. In general, the four human influence datasets omitted all linear disturbance types much more than the intactness datasets, 49-74% on average compared to 0-8% for the intactness datasets. The VLIA2015 map omitted all disturbance types more often than any other map. The HFP2013 map was the only human influence dataset that had some omission rates below 20%, specifically for railways and dams. Among the human influence datasets, omission of seismic lines ranged from 62% by the AB2015 map to 89% by the VLIA2015 map. This compares to 2-7% for the intactness datasets. Roads, pipelines, and powerlines were also omitted 37-81% of the time by the human influence datasets compared to 0-8% for the intactness datasets. Among the intactness group, the CIFL2013 and GIFL2016 performed best, omitting only 2-4% of seismic lines and roads, respectively. Only dams were omitted more than 10% of the time, by the HA2010 and CIFL2013 maps. The GIFL2016 map never exceed an omission rate of 5% while the HA2010 consistently omitted more linear disturbances than CIFL2013 and GIFL2016.

By far, the most common and widely distributed anthropogenic polygonal disturbances in the study area were forest cutblocks followed by agriculture, settlements, and mines. Well sites and other oil and gas infrastructure also occurred to a much lesser extent. In general, and as with linear disturbances, the four human influence datasets omitted all polygonal disturbance types more than the intactness datasets, 10-73% on average compared to 0.2-2% for the intactness datasets. In particular, cutblocks were omitted 57-91% of the time by the human influence datasets. This compares to 0.1-5% for the intactness datasets. Mines and well sites were also more often omitted by the human influence maps. Among the human influence datasets, the HFP2013 map had the lowest omission rate for all polygonal disturbance types except for well sites. Both CIFL2013 and GIFL2013 never exceeded a 1% omission rate for any polygonal disturbance type. Similar to the linear disturbances, the HA2010 map had higher omission rates than the two IFL maps, although it never exceeded 5%.

The use of higher resolution test data was most noticeable with linear disturbances, with 68% and 28% seismic lines and roads identified, respectively. This had much larger impact on the intactness datasets, roughly doubling the omission rates of roads for HA2010, CIFL2013 and GIFL2013 and of seismic lines for HA2010 and CIFL2013. Even with these increases, however, the overall omission rates for all linear disturbances for CIFL2013 and GIFL2013 was less than 6.5%. In contrast, the omission rates for the human influence datasets did not change much but remained much higher than for the intactness datasets.

### Case studies

The supporting information includes results from two case studies. The first case study evaluates the effectiveness of the intactness and human influence maps at identifying disturbances related to placer mining in west-central Yukon (S2_case_study_1.html). Among the 7 datasets we analysed, the GIFL2016 map omitted the least amount of both linear and polygonal anthropogenic disturbances (0.0% and 1.9%, respectively; Table 1). However, it only identified only 12.3% of the study region as being intact, far less than the CIFL2013 and HFP2013 maps which identified 49% and 55% of the area as intact, respectively. Three of the datasets, VLIA2015, GHM2016 and AB2015, identified 92-96% of the area as being intact and, consequently had very high rates of omission, ranging from 66-95% for linear disturbances and 88-98% for polygonal disturbances. The other two datasets, CIFL2013 and HA2010 omitted a moderate amount of both polygonal (12% and 22%, respectively) and linear (21% and 38%, respectively) disturbances. The second case study consists of a simple two factor analysis conducted in northern Alberta to illustrate the sensitivity of intactness estimates to buffer size and minimum intact patch size (S3_case_study_2.html). The results indicate that the area estimated to be intact varies from 44-98% depending of buffer size and minimum intact patch area, with buffer size having a larger influence than minimum patch size (Table S3.2, Figure S3.2).

## Discussion

The boreal region of North America is experiencing rapid industrial development (Brandt et al. 2013, Venier et al. 2014, White et al. 2017). Consequently, there is a need for reliable and up-to-date information on changes in ecosystem conditions. To that end, we compared eight global and regional maps depicting intactness or cumulative human influence on ecosystems in the boreal region. Our results revealed large differences in the area estimated to be intact or relatively free from human influence. In Canada, estimates ranged from 55-89% while in Alaska they ranged even more, from 32-96%. Likewise, the similarity between pairs of datasets in the Canadian boreal ranged from 0.58 to 0.86 on a scale of 0-1. In total, 45% of the region was identified as intact by the seven most recent datasets. This variation was also evident in the ability of the datasets to account for anthropogenic disturbances that are increasingly common in the boreal region, especially those associated with resource extraction. The four human influence datasets (Human Footprint, Global Human Modification, Large Intact Areas, and Anthromes), in particular, omitted 59-85% of all linear disturbances and 54-89% of all polygonal disturbances. In contrast, the Global IFL, Canada IFL, and Human Access maps omitted 2-7% of linear disturbances and 0.1-5% of polygonal disturbances. Several differences in map characteristics, including input datasets and methods used to develop the maps may help explain those differences.

Input datasets appear to play an important role in the observed variation in intactness estimates, spatial agreement between maps, and the ability to detect both linear and polygonal anthropogenic disturbances. Among the datasets evaluated, the four human influence maps relied mostly on combining existing thematic maps that each represented one stressor into a cumulative disturbance map. The primary stressors used were mostly related to settlement, agriculture, population density and transportation, with little information on resource extraction activities. Exceptions included the use of a forest cover change map (Hansen et al. 2013) by the Low Impact Areas dataset and mining and oil wells by the Global Human Modification dataset. Even so, this did not make a big difference in the omission rates of those disturbances. Moreover, the majority of input datasets were raster maps with a resolution that was generally ≥1-km^2^, which also contributed to the omission of finer-scale anthropogenic changes and disturbances. In contrast, the Human Access and IFL maps mostly relied on processing high resolution satellite imagery (i.e., 30-m) along with some thematic maps to identify disturbances. This resulted in far fewer omissions of anthropogenic disturbances related to resource development such as forest cutblocks, pipelines, seismic lines and roads. However, the use of even finer resolution test data (i.e., 15-m) revealed increased omission rates, especially for seismic cutlines and roads, whose width make them particularly challenging to detect without imagery of an appropriate resolution. In the case of seismic lines, there has also been a reduction in their width over time which would also contribute to newer lines being undetected by satellite imagery (Lee and Boutin 2006, van Rensen et al. 2015).

The age and temporal resolution (i.e., how often they are updated) of datasets also has important implications for their suitability for conservation planning, especially in areas of the boreal that are rapidly changing, including the boreal plains of western Canada, and southern parts of the boreal shield in Ontario and Quebec (Government of Quebec 2009, OMNR 2013). Older datasets that were only produced once, such as Frontier Forests, may be useful from a historical perspective but would be a poor choice for conservation planning. More recent datasets, such as the Human Access and Canada IFL maps, have now been discontinued leaving only the Global IFL map as a true intactness dataset. The increasingly rapid pace of industrial development means that even recently produced intactness maps are quickly out-of-date, suggesting the importance of updating maps on a regular basis, ideally annually. Three datasets, Human Footprint, Anthromes and Global IFL, stood out for having at least three updated products between 2000-2016, allowing for monitoring and change detection based on consistent and replicable methods. Newer datasets, such as the Global Human Modification and Low Impact Areas, have only one temporal product but may provide a complementary approach to the Human Footprint for assessing ecosystem conditions at a global scale. In fact, despite their differences, those three datasets along with the Anthromes datasets provided similar estimates of the amount of remaining terrestrial ecosystems with very low human influence (Riggio et al. 2020).

Methodological differences among maps were mostly related to study area delineation, minimum intact patch size, and the use of exclusion buffers around linear and polygonal anthropogenic disturbances. For example, some of the discrepancies between the Frontier Forests and Canada IFL maps are due to the delineation of the Frontier Forests forest zone, which excluded northern, less densely forested portions of the Canadian boreal. Similarly, the Global IFL maps used a satellite-based global tree cover map to define their study area, resulting in some parts of the boreal region being excluded because tree canopy was < 20%. The use of a minimum intact patch size also contributed to discrepancies among maps, with four of the maps specifying a minimum size. The Global IFL maps, for example, considered that an intact forest should have a minimum size of 50,000 ha (Potapov et al. 2017). In contrast, the Canada IFL maps used a minimum threshold of 5,000 ha for boreal ecozones and 1,000 ha for temperate ecozones (Smith and Cheng 2016); the latter only occurred along the southern edge of the boreal region. Consequently, a greater total area of intact forests was identified by the Canada IFL maps. Other maps, such as the Human Access map, did not have a minimum area requirement and consequently identified an even greater amount of intact area. This resulted in higher omission rates for linear and polygonal disturbances in comparison to the IFL maps, in particular in areas identified as being intact and smaller than the minimum patch size used by the other maps.

The applied widths of human influence zones (or buffers) also contributed to differences in the extent of mapped intact areas. For example, the Human Footprint maps considered up to 15-km wide zones of influence around features such as roads, major rivers and coastlines, since they are often used as transportation corridors or have high population densities. While there is plenty of evidence that human activities can have impacts beyond the point source (e.g., wolf avoidance of areas with human activities (Shepherd and Whittington 2006); impacts of riparian forest harvesting on streams (Richardson and Béraud 2014)), the use of thresholds eliminated many areas considered intact by the Human Access and IFL maps. This may be justified in some coastal zones of Europe and more populated regions of North America, but it is not as well supported in remote areas of the northern boreal forest, where population density is negligible. Our second case study provided a simple illustration of the sensitivity of IFL estimates to the size of exclusion buffers and the minimum intact patch size on intactness estimates (Supp Info). In particular, the use of buffer exclusion zones by themselves resulted in a much greater reduction in intact areas than the use of a minimum intact patch size criteria on its own. The use of simple buffers around disturbances limits the users’ ability to use a more flexible and nuanced approach to allocating degrees of intactness within areas that have not been disturbed but are close to a disturbance. For example, when identifying reserves for species that have strong avoidance of human-impacted areas such as caribou (Environment Canada 2008), these buffers may be appropriate, and would not represent an underestimation of intact areas. However, when conservation efforts focus on less sensitive species, these buffers may be too conservative and underestimate the amount of suitable habitat. To be most flexible, intactness mapping projects could avoid using buffers or at least provide underlying unbuffered data.

Overall, and as with input data, the datasets we evaluated can be broadly divided into two groups based on similarities in their methodology, with the IFL and Human Access maps belonging to one group and the four human influence maps belonging to the other. However, even within groups, minor differences in methods resulted in relatively important differences in the areas identified as intact. For example, the Global IFL maps considered all wildfires occurring in proximity to infrastructure (e.g., settlements) as non-intact, resulting in less intact area identified in comparison to the Canada IFL maps (Lee 2009). Fires play crucial roles in the dynamics of Canadian boreal forests, where most of the area burned is due to lightning-caused fires (Price et al. 2013). This alone would account for an under-estimation of 400,000 km^2^ of intact boreal and temperate forests in Canada by the Global IFL maps (Lee 2009). Another source of disagreement was due to the treatment of rivers affected by hydroelectric power generation, which were excluded using a 1-km buffer by the Global IFL maps but not by the Canada IFL and Human Access maps.

Global maps such as the Anthropogenic Biomes, Global Human Modification, Low Impact Areas and Human Footprint maps may be appropriate for broad-scale conservation assessments where finer resolution data are not available. For example, this approach was used to identify and prioritize global wilderness areas (Mittermeier et al. 2003), identify remaining intact areas globally and within biomes (Riggio et al. 2020), and analyse the connectivity of protected areas via intact land (Ward et al. 2020). However, obtaining more detailed and up-to-date regional maps of intactness or disturbances should be a priority for any systematic conservation planning exercise, in the boreal or elsewhere. Increasingly, researchers are using freely available satellite imagery to produce time series of high resolution land cover maps, including maps that track changes in forest disturbances (e.g., White et al. 2017). In addition, there exists several examples of regional intactness and human influence maps in North America and other parts of the world. For example, the Human Footprint approach has been applied at regional scales in the United States and Canada (Woolmer et al. 2008, Leu et al. 2008). Other recent related initiatives have aimed to characterize landscape patterns, forest fragmentation, and forest change at regional (Raiter et al. 2017), national (Wulder et al. 2008, Pasher et al. 2013, Guindon et al. 2014, White et al. 2017) and global scales (Hansen et al. 2013). Regional datasets also afford greater sophistication by integrating information on context, connectivity, habitat, and species (e.g., Plumptre et al. 2019, Grantham et al. 2020, Mokany et al. 2020). As an indication of the importance of considering intact areas in conservation planning, a major international conference on “Intact Forests in the 21st Century” recently took place in Oxford in 2018^1^ to discuss regional and global approaches.

The development of intactness and human influence datasets has also led to some critiques on the utility of the concept of intactness (e.g., Innes and Er 2002, Bernier et al. 2017, Venier et al. 2018) and debates amongst mapping methods (e.g., Kennedy et al. 2020, Venter et al. 2020, Riggio et al. 2020). Two recent papers, with relevance to the boreal context, argue for a more sophisticated approach to the assessment of the loss of ecological value from forests. Bernier et al. (2017) reviewed the concept of “primary forest” as a metric of forest environmental quality, and its use by the Food and Agriculture Organization (FAO) for reporting country-level statistics. Of particular concern is the lack of a consistent operational definition resulting in substantial differences in the way primary forest areas are defined and measured within each country. They note that more recent approaches, such as IFLs, provide more consistency by using satellite imagery but do not consider regional differences in ecosystem processes that can result in large differences in areas identified as intact. Venier et al. (2018) distinguished between conceptual and operational definitions of an IFL and provides a historical review of the intact forest landscape concept and intactness mapping, both globally and regionally. Both papers point out limitations in the criteria used to map intact areas and argue for a more sophisticated approach, one that considers intactness as a gradient rather than a binary condition and where the minimum patch size is not standardized but guided by regional ecological conditions and processes. Specifically, the standard operational definition of a Global IFL sets a minimum intact patch size of 50,000 ha, which is arbitrary and disconnected from regional ecosystem processes which may require a smaller or larger minimum size. For example, the minimum intact patch size may be too small for wide ranging species such as caribou and wolverine and for ecosystem processes such as wildfires which can exceed 1,000,000 ha in the boreal region. Ideally, the minimum size that an intact patch needs for conservation planning should be related to habitat requirements for focal species and ecological processes (Haddad et al. 2015).

Our analysis has some limitations. For example, we only reviewed existing datasets that were freely available and covered the boreal region of Canada at a minimum, but we did not consider regional datasets. An evaluation of regional intactness and human influence datasets would be useful since many conservation decisions are made at those scales. In addition, our analysis represents a snapshot in time as i) datasets are continually being produced, revised or updated and ii) anthropogenic disturbances are happening every day in the boreal region. Our evaluation of both intactness and human influence datasets also required that some datasets be reclassified to binary maps identifying only areas with minimum human impacts. However, not all datasets had a clear “no impact” class, and consequently, we reclassified some of the maps to create an analogous class showing areas with little or no influence. For one dataset, the Global Human Modification map, we used values ranging from 0-0.01 to be consistent with methods used by Riggio et al. (2020).

The boreal region of North America is currently undergoing rapid industrial development, and there is an urgent need to quantify changes in ecosystem conditions and to identify new protected areas. Several datasets have been developed over the last two decades that can be used to assist in monitoring, conservation planning, and adaptive management. The datasets can be broadly grouped into i) those that combine existing stressor datasets to create a cumulative human influence map and ii) those that process satellite imagery to identify and map disturbances. For all linear and polygon anthropogenic disturbance types, the former group was shown to be far less effective than the latter at incorporating anthropogenic disturbances related to resource development in the boreal. However, even among the latter group, the use of buffers and minimum patch sizes limit the flexibility of those products to enable more sophisticated conservation planning. Encouragingly, the increasing concern due to climate and land use change is leading to the continued refinement or revision of existing datasets over time and the development of new products using more sophisticated approaches and finer resolution input data. Moreover, many datasets are being provided freely and some with accompanying algorithms and code used to develop them (e.g., the Human Footprint), allowing approaches to be replicated or adapted for regional use. In addition, many Canadian provinces are now making available historical and modern datasets related to resource management and development. Ultimately, the decision on which dataset to use will depend on the objectives of the conservation planning initiative and the availability of the most recent high quality datasets available in the planning region. Our goal was to contribute to the review and assessment of some widely available datasets and provide some guidance for their use in the boreal context.

## Supporting information

The following supplementary tables and maps are available on GitHub (https://github.com/prvernier/intactness):

- S1_datasets.md. Description of intactness and human influence datasets within the boreal region of Canada and Alaska.
- S2_case_study_1.html. Effectiveness of intactness and human influence maps at identifying disturbances related to placer mining in west-central Yukon.
- S3_case_study_2.html. Sensitivity of intactness estimates to buffer size and minimum patch size.

## Footnotes

1 https://www.eci.ox.ac.uk/if21/

